# LOCOM2: Robust Differential Abundance Analysis for Microbiome Data

**DOI:** 10.64898/2026.04.07.716976

**Authors:** Mengyu He, Glen A. Satten, Yi-Juan Hu

## Abstract

**Background:** Numerous methods have been developed for differential abundance analysis of microbiome data; however, many fail to adequately control error rates, contributing to the reproducibility crisis in microbiome research. Moreover, new challenges have emerged, including large-scale studies, differential library size distributions, unbalanced case–control designs, and the increasing availability of only relative-abundance data rather than read counts.

**Methods:** We propose LOCOM2 to address these challenges. The method refines the weighting scheme in LOCOM to eliminate confounding by library size while accommodating relative abundance data. It incorporates a series of adjustments to ensure stable and reliable estimation, even under extreme conditions such as very rare taxa and highly unbalanced case–control designs. In addition, LOCOM2 replaces the computationally intensive permutation procedure in LOCOM with a Wald-type test, substantially improving computational efficiency. To evaluate performance, we conducted extensive simulation studies using the MIDASim simulator and three data templates representing diverse body sites. We benchmarked LOCOM2 against state-of-the-art methods, including LOCOM, LinDA, ANCOM-BC2, MaAsLin2, and MaAsLin3. This benchmarking effort provides an essential foundation for the next stage of microbiome research.

**Results:** LOCOM2 achieved accurate control of the false discovery rate across all simulation scenarios, whereas none of the other methods consistently did so. LOCOM2 also demonstrated the highest sensitivity for detecting true signals. Applications of these methods to three real microbiome datasets further corroborated these findings.

## Background

There is growing evidence that the human microbiome plays a critical role in human health and diseases. Beyond its well-known functions in digestion and nutrition, the microbiome also engages in systemic communication through axes such as the gut–brain [1] and gut–heart axes [2].

Despite these advances, microbiome research faces a profound reproducibility crisis, largely attributable to the complexity of data. Sequencing-based microbiome data are compositional (a change in one taxon’s relative abundance necessarily alters others due to the constant-sum constraint), sparse (often containing 80–90% zeros), overdispersed (reflecting substantial interindividual heterogeneity), and highly susceptible to experimental bias [3]. These complexities can easily invalidate statistical analyses if not properly addressed. Indeed, [4] reported that 14 differential abundance methods produced divergent results across 38 datasets. Studies of differential abundance methods [5, 6, 7, 8, 9, 10, 11] have repeatedly shown that most existing methods fail to reliably control error rates, which may substantially contribute to the lack of reproducibility. Therefore, future progress in microbiome research hinges on the development and application of statistical methods that rigorously address these data complexities.

Despite these challenges, differential abundance analysis is routinely applied to microbiome research, to attempt to identify microbial biomarkers or therapeutic targets. The numerous methods for microbiome differential abundance analysis can be broadly grouped into two categories. The first category, compositional methods, includes LOCOM [12], LinDA [13], and ANCOM-BC2 [14], which account for the compositional constraint through log-ratio transformations or via bias correction. Compositional methods have the potential to produce bias-free findings, although this depends on correct handling of zero counts [15]. The second category, non-compositional methods, includes LEfSe [16] and MaAsLin2 [5], which treat taxa independently without accounting for the compositional constraint. Recently, LEfSe and MaAsLin2 have been superseded by MaAsLin3 [17], which explicitly accounts for compositionality. To address the high prevalence of zeros, LinDA, ANCOM-BC2, and MaAsLin2 employ *ad hoc* pseudo-counts or imputations, which lack a solid statistical foundation and are not robust to experimental bias [3, 15]. MaAsLin3 [17] models nonzero data and zero data separately (i.e., applying a log-linear regression to non-zero data and logistic regression to presence-absence data) and then combining the results, thus ignoring the sampling nature of many zeros and increasing sensitivity to library size variation. LOCOM treats zeros as natural observations rather than missing values, thereby preserving statistical rigor. Figure 1 illustrates the use of these methods in clinical research over the past three years, showing that LEfSe and MaAsLin dominate in practice. The relatively low adoption of LOCOM may stem from its installation difficulties, absence from CRAN (the standard repository of R packages), and relatively long runtime.

**Figure 1:**
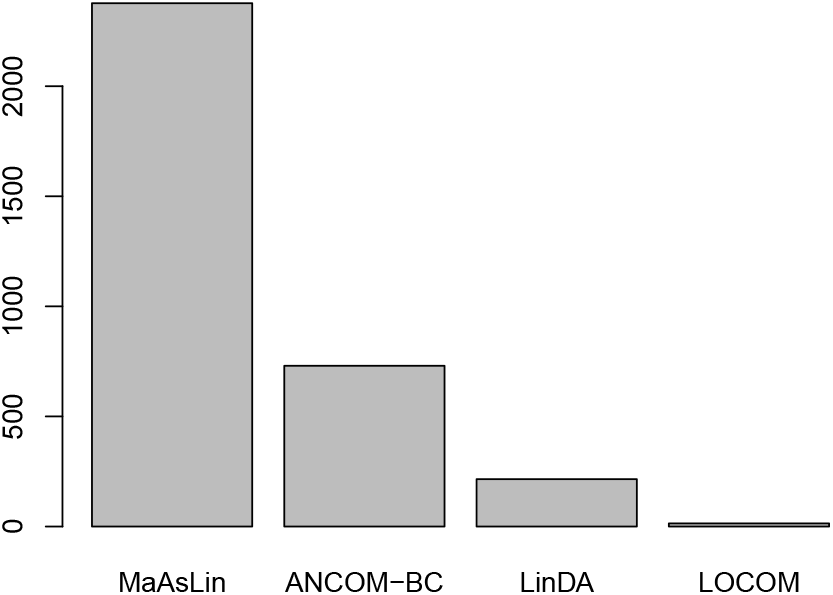
Method usage between 2022 and 2025. “MaAsLin” refers collectively to MaAsLin [46], MaAsLin2, MaAsLin3, and LEfSe, as these methods were developed by the same research group. “ANCOM-BC” refers to both ANCOM-BC [47] and ANCOM-BC2.

New challenges have recently emerged in differential abundance analysis. Large-scale microbiome studies are becoming increasingly common. For example, a study of Canadian population [18] included 1,190 individuals; the Finnish National FINRISK study [19] included 5,959 individuals; and a study of German cohorts [20] included 8,956 individuals. In such large-scale studies, samples are inevitably sequenced across multiple batches, often leading to differential library size distributions between case and control groups [14]. In addition, unbalanced two-group design is gaining prevalence [21], particularly in large-scale observational studies or when analyses extend to secondary traits. Moreover, shotgun metagenomic sequencing is rapidly expanding as an alternative to 16S rRNA gene sequencing. Commonly used bioinformatics pipelines for metagenomic data, such as MetaPhlAn [22], Kraken [23], and HUMAnN [24], output taxon relative abundances directly rather than read counts, as read counts assigned to a species depends on its genome size.

LOCOM may encounter difficulties under these conditions. Because it models read count data using logistic regression, it is sensitive to differential library sizes and is not applicable to relative abundance data. Its reliance on permutation-based inference limits scalability in large studies. In addition, as it is not robust to rare taxa, it requires stringent filtering to exclude them and is not well suited to highly unbalanced case–control designs. Nevertheless, given LOCOM’s strong statistical foundation for addressing compositionality and zeros, extending LOCOM to overcome these limitations is highly desirable.

Tools for evaluating performance of differential abundance methods have advanced considerably in recent years. MIDASim [25] is a state-of-the-art simulator for microbiome data, generating replicates that preserve key features (e.g., relative abundance distributions, zero proportions, and overdispersion) of a template microbiome dataset while allowing incorporation of covariate effects. By contrast, other microbiome data simulators, such as the Dirichlet–Multinomial model [26], metaSPARSim [27], SparseDOSSA [28], and MB-GAN [29], either fail to adequately capture these essential data features or require prohibitive computational resources [25]. Notably, recent benchmarking studies [9, 10, 11] have relied on these suboptimal simulators. Therefore, a benchmarking study of differential abundance methods that leverages MIDASim for rigorous evaluation, particularly under the newly emerging conditions not previously examined (e.g., large-scale studies), remains lacking.

To fill this gap, this paper has two objectives. First, we introduce LOCOM2 to address the aforementioned limitations, making it applicable to relative abundance data, computationally efficient for large datasets, and robust to differential library sizes, unbalanced case–control designs, and rare taxa. Second, we conduct MIDAsim-based simulation studies to benchmark LOCOM2 against LOCOM and other state-of-the-art methods. Table 1 summarizes the methods evaluated in this study along with their key features. LEfSe was excluded because it does not account for compositionality and cannot adjust for additional covariates.

**Table 1:** Differential abundant analysis methods evaluated in this study.

| Method | Addressing<br>compositional<br>effects | Bypassing<br>zero<br>imputation | Supporting<br>RA only<br>data | Scaling<br>to large-scale<br>data |
| --- | --- | --- | --- | --- |
| LOCOM2 | ✓ | ✓ | ✓ | ✓ |
| LOCOM2-P | ✓ | ✓ | ✓ | × |
| LOCOM | ✓ | ✓ | × | × |
| LinDA | ✓ | × | ✓ | ✓ |
| ANCOM-BC2 | ✓ | × | × | ✓ |
| MaAsLin2 | × | × | ✓ | ✓ |
| MaAsLin3 | ✓ | ✓ | ✓ | ✓ |

Note: RA=relative abundance. “✓” and “×” indicate “Yes” and “No”, respectively.

## Methods

### Review of LOCOM

Let *X*_*ij*_ denote the observed read count for taxon *j* in sample *i*, where *j* = 1, …, *J* and *i* = 1, …, *n*. Define the *observed* relative abundance as 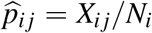, where *N*_*i*_ = ∑_*j*_′ *X*_*i j*_′ is the library size. Studies of model communities [30] have shown that, because of experimental bias, the expectation of 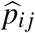, denoted as *p*_*ij*_, is linked to the *true* underlying relative abundance *π*_*ij*_ through log (*p*_*ij*_) = *γ* _*j*_ + log(*π*_*ij*_) + *α*_*i*_, where *γ* _*j*_ is a taxon-specific bias factor describing how experimental bias distorts the observed relative abundance, and *α*_*i*_ is a normalization factor ensuring 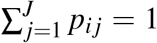. Let *Z*_*i*_ denote the covariate vector, which usually includes the trait of interest *Y*_*i*_ and additional covariates *C*_*i*_, so that *Z*_*i*_ = (*Y*_*i*_,*C*_*i*_). We further assume that *Z*_*i*_ is related to *π*_*ij*_ via 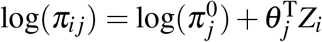, where *θ*_*j*_ is the association parameter and 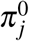 denotes the baseline relative abundance (when *Z*_*i*_ = 0). This yields a set of (*J* − 1) equations:

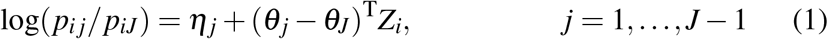

where, without loss of generality, *J* denotes the reference taxon, whose choice does not affect the final result as it cancels out in later derivations. Each 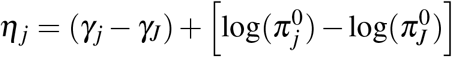 is treated as a nuisance parameter. Because bias factors only affect the *η*_*j*_s, inference on the *θ*_*j*_s is fully robust to experimental bias. To ensure identifiability, we set *θ*_*J*_ = 0. It is worth noting that the log odds ratios *θ*_*j*_ correspond to additive-log-ratio transformations applied to the parameters *p*_*ij*_, rather than the observed data. This formulation allows LOCOM to operate directly on the original count data without requiring zero imputation, which constitutes a key advantage of the method.

When the number of taxa *J* is large, jointly fitting the polytomous logistic regression in (1) can be numerically challenging. Instead, following [31], LOCOM approximates the solution by fitting a series of dichotomous logistic regressions, each comparing a taxon to the reference taxon (practically chosen to be the taxon with the highest mean relative abundance). Define the relative abundance of taxon *j* in the subcomposition comprising only taxon *j* and the reference taxon *J* as *µ*_*ij*_ = *p*_*ij*_*/*(*p*_*ij*_ + *p*_*iJ*_). Equation (1) implies a logistic regression model for each taxon *j* (*j* ≠ *J*) against taxon *J*:

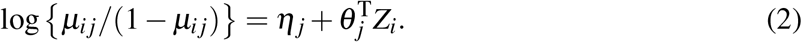

Thus, LOCOM fits separate logistic regressions to estimate each *θ*_*j*_. To account for the possibility of separation when all counts of a taxon are zero in either the case or control group [32], LOCOM uses the Firth-corrected likelihood [33]. Specifically, LOCOM estimates *θ*_*j*_ by solving the Firthcorrected score equation:

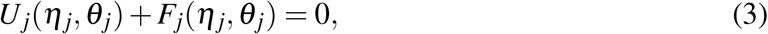

where

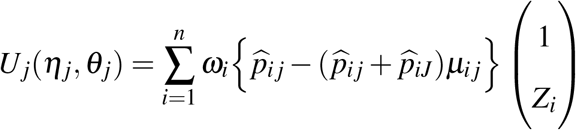

is the score function, *ω*_*i*_ = *N*_*i*_ weights each observation in proportion to its library size, and *F*_*j*_(*η*_*j*_, *θ*_*j*_) is the Firth correction to the score function (see [12]).

We typically write *θ*_*j*_ = (*β*_*j*_, *ϕ*_*j*_), where *β*_*j*_ corresponds to the trait of interest *Y*_*i*_ and thus the parameter we wish to test and *ϕ*_*j*_ corresponds to the additional covariates *C*_*i*_ that we need to control for confounding. The resulting estimator of *β*_*j*_ is denoted by 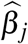.

To test whether taxon *j* (*j* = 1, …, *J*) is associated with the trait, LOCOM uses the test statistic 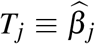 − median 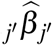, where median _*j*_′ denotes the median over all taxa *j*^*′*^ = 1, …, *J*. Centering 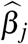 by the median removes dependence on the choice of reference taxon. Under the assumption that more than half of taxa are null, the median coefficient corresponds to a null taxon, regardless of whether the reference taxon *J* is null. Consequently, a null taxon *j* satisfies *β*_*j*_ − median_*j′*=1,…,*J*_*β*_*j*_′ ≈ 0, rather than simply *β*_*j*_ = 0.

LOCOM employs permutation-based inference to circumvent the difficulty of estimating the variance of *T*_*j*_. If there are no additional covariates, LOCOM permutes the values of *Z* ≡ *Y*. When additional covariates are present, LOCOM follows a residual permutation scheme proposed by Potter [34]. First, we regress *Y* on *C* to obtain the residuals *Y*_*ε*_, and replace *Z* by 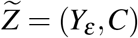; note that fitting (2) using 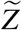 in place of *Z* does not affect the estimate 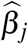. For each permutation *r* (1 ≤ *r* ≤ *R*), we permute 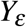 to obtain 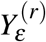 and define 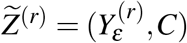. We then refit the model using 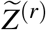 for each taxon *j* to obtain 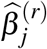. Inference is then based on the distribution of 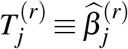−median 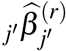. Although Sandve’s sequential stopping rule [35] is applied to terminate permutations adaptively and as early as possible, when some true *p*-values are close to the significance threshold, the procedure may still require a very large number of replicates (often *>* 50, 000) before termination.

### LOCOM2 adopts new weights

LOCOM uses weights *ω*_*i*_ = *N*_*i*_ in (3), so each sample is weighted by its library size, giving greater influence to samples with larger libraries in parameter estimation; this weighting is optimal when the magnitude of the raw counts is informative. However, library size is a technical artifact rather than a biological quantity, and therefore does not reflect the importance of an observation. Further, when library size distributions differ between case and control groups, such weighting may introduce confounding and lead to spurious findings.

LOCOM2 uses uniform weights for all samples, i.e., *ω*_*i*_ = 1 for all *i* in (3). This weighting scheme eliminates potential confounding by library size by ensuring that all samples contribute equally. An additional advantage is that it enables direct analysis of relative abundance data without requiring access to raw counts. This is particularly important for combining data from multiple studies, where relative abundances rather than raw counts are most frequently shared.

When *ω*_*i*_ = 1, the *U*_*j*_(*η*_*j*_, *θ*_*j*_) term in equation (3) is no longer the score function of a likelihood, but instead defines a generalized estimating equation (GEE). We therefore modify *F*_*j*_(*η*_*j*_, *θ*_*j*_) in (3) to incorporate the bias-reduction adjustment for GEEs developed by [36]. We also follow their recommendation to add a Jeffreys-type penalty, ensuring that the bias-reduced estimates are always finite [37]. Careful attention to the overall scale must be paid to ensure that the adjustment terms are of higher order than the leading term *U*_*j*_(*η*_*j*_, *θ*_*j*_) in (3). Further details and the final form of the adjusted estimating equation are provided in the Supplementary Materials. Together, these techniques enhance LOCOM2’s ability to handle rare taxa and unbalanced case-control data.

### LOCOM2 employs Wald-type tests

Permutation-based inference, as implemented in LOCOM, is computationally expensive and does not scale well to large studies, as a large number of permutations is required to obtain sufficiently small *p*-values needed for significance when testing many hypotheses. In contrast, the analytical variance estimator for the test statistic *T*_*j*_, which takes a sandwich form derived from GEE theory [38] (ignoring the higher-order variance contribution from the median adjustment), does not perform well for rare taxa, even in large studies (results not shown).

LOCOM2 retains the residual-permutation scheme of Potter [34] used in LOCOM to generate permutation replicates. However, LOCOM2 adopts a simplified testing strategy in which a modest number of permutation (null) replicates (e.g., *R* = 1000) is used to estimate a variance-covariance matrix that is then used in a pseudo-Wald test. This approach is motivated by our observation that, for null taxa, each test statistic *T*_*j*_ is approximately normally distributed across simulation replicates (Figures S1–S2). Further, we also find that the empirical variance of 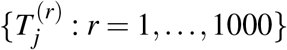 from a single simulated dataset agrees with the empirical variance of *T*_*j*_ calculated using 1000 simulated datasets (Figure S3). While these findings support the proposed approach, we notice from Figure S4 that the normal approximation is less accurate for rare taxa, particularly for unbalanced case–control data. To improve data normality, we apply a taxon-specific Yeo-Johnson (Y-J) transformation [39] to 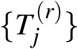 using the `yeojohnson` function in the R package `bestNormalize`, obtaining transformed statistics 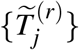. Using the same Y-J transformation parameter 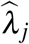, we transform the observed statistic *T*_*j*_ to obtain 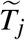. These values are then used to construct the pseudo-Wald statistic:

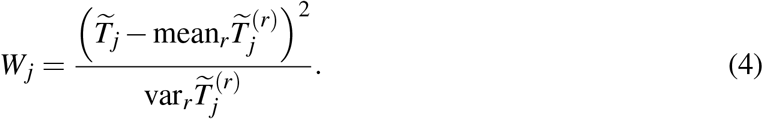

where mean_*r*_ and var_*r*_ denote the mean and variance across permutation replicates. The *p*-value is obtained by comparing *W*_*j*_ to a chi-squared distribution with one degree of freedom under the null. Because LOCOM2 relies on this asymptotic approximation, only a modest number (*R* = 1000) of permutations is required.

### LOCOM2 filter

Very rare taxa, those with few non-zero observations, should be filtered out, as they are often unreliable (e.g., due to artifacts), contribute little statistical information, inflate the multiple-testing burden, and can violate modeling assumptions, thereby reducing power and stability. LOCOM requires filtering out taxa present in fewer than 20% of samples, while LinDA, ANCOM-BC2, and MaAsLin2 apply a default filter that excludes taxa present in fewer than 10% of samples. (MaAsLin3 does not apply filtering by default, but recommends some, albeit unspecified, filtering to improve statistical power.) Given the sparsity of microbiome data, these criteria can remove too many taxa and may miss important signals. Moreover, as sample size increases, these criteria become increasingly stringent in terms of the number of samples that a taxon must appear in, turning increasing sample size into a liability rather than an advantage.

Because LOCOM2 has improved capabilities for handling rare taxa, we introduce a new filtering rule that retains a taxon if it is present in at least 10% of samples or in at least 10 samples, whichever threshold is lower. This rule reduces to the LinDA/ANCOM-BC2/MaAsLin2 filter for studies with 100 or fewer samples. Importantly, it avoids penalizing larger datasets as sample size increases.

## Results

### Simulation studies

Using the MIDASim simulator [25], we conducted extensive simulation studies based on three data templates representing the major body sites in microbiome research: 1) an upper-respiratory-tract (URT) microbiome dataset [40] (57 samples, 856 taxa), 2) a gut microbiome dataset from a Crohn’s disease (CD) cohort [41] (109 samples, 2,529 taxa), and 3) a vaginal microbiome dataset from the MOMS-PI cohort [42] (517 baseline samples, 1,146 taxa). As shown in Figure 2, the proportion of rare taxa varies across the three body sites, being lowest in the URT and highest in the vaginal site. For each template, we generated replicate datasets with sample sizes of *n* = 100, 1, 000, and 10, 000.

**Figure 2:**
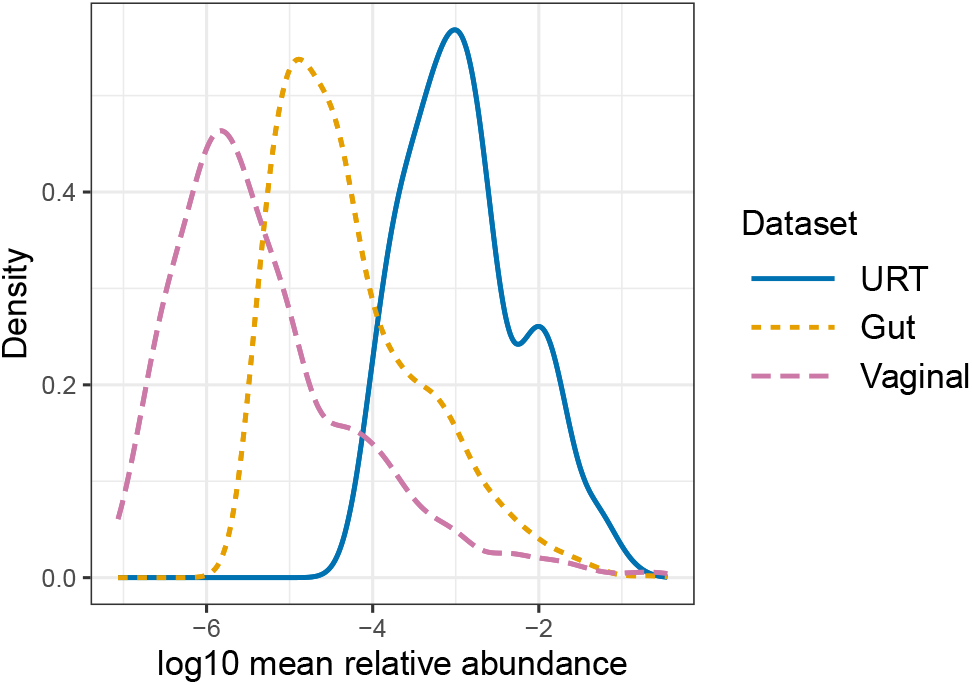
Distributions of mean relative abundances (log10 scale) of taxa in the URT, gut (CD cohort) and vaginal microbiome datasets.

We simulated either a binary or continuous trait *Y*_*i*_ for each sample; for some simulations we also generated an additional covariate *C*_*i*_. For the binary trait, corresponding to disease status, we first considered the most common scenario with an equal (50:50) case:control ratio, and then extended the analysis to increasingly unbalanced ratios of 30:70, 70:30, 10:90, and 90:10. In simulations based on the URT microbiome, 20 taxa with mean relative abundances greater than 0.2% were randomly selected (from the original data) as associated with *Y*_*i*_. The covariate *C*_*i*_ was binary, assigned so that within each trait group, half of the samples had *C*_*i*_ = 0 and the other half had *C*_*i*_ = 1. In simulations based on the gut microbiome, the associated taxa were the 106 taxa identified as differentially abundant in the original data. In this setting, *C*_*i*_ was generated by randomly sampling from the observed age values, and was treated as a continuous variable. In simulations based on the vaginal microbiome, 20 taxa with mean relative abundances greater than 0.05% were randomly selected as associated with *Y*_*i*_, and no covariate was included. For continuous-trait simulations, *Y*_*i*_ was drawn from a uniform distribution on [−1, 1]. The covariate *C*_*i*_ was binary (0/1), with half of the samples assigned *C*_*i*_ = 1 in simulations based on the URT data, and was continuous (randomly sampled age values) in simulations based on the gut data.

MIDASim incorporates the effects of *Y*_*i*_ and *C*_*i*_ into the relative abundance *π*_*ij*_ through the following multiplicative model:

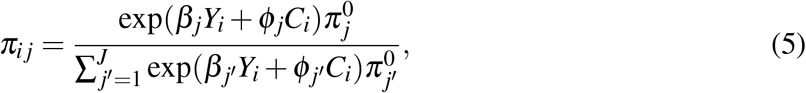

where *β*_*j*_ and *ϕ*_*j*_ denote the effect sizes of *T*_*i*_ and *C*_*i*_ on taxon *j*, respectively, and 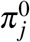 is the baseline relative abundance of taxon *j* in the null data template. The URT and vaginal datasets were used directly as null templates. For the gut dataset (from the CD cohort), the null template was constructed by removing the effects of CD and age at their associated taxa, as these effects were strong. Notably, model (5) reflects the core assumption underlying all compositional methods. Aside from this point, we believe the MIDASim-based simulations do not preferentially favor LOCOM or LOCOM2. In the URT and vaginal microbiome simulations, we set *β*_*j*_ = 0 for null taxa and *β*_*j*_ = *β* for taxa associated with the trait *Y*_*i*_, assuming a common effect size; *β* was varied to assess sensitivity. We set *ϕ*_*j*_ = 1 for taxa associated with the covariate *C*_*i*_ and *ϕ*_*j*_ = 0 otherwise. In the gut microbiome simulations, we specified 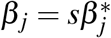, where 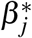 denotes the estimated effect size (which may be positive or negative) for taxa identified as associated with CD in the real data, and *s* is a scaling factor varied to assess sensitivity; we set *β*_*j*_ = 0 for null taxa. Similarly, *ϕ*_*j*_ was set to the estimated effect size for taxa associated with age and to 0 otherwise. Because the estimated effect sizes 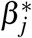 varied in direction and were largely balanced (Figure S5), the resulting compositional effects in the gut microbiome simulations were much smaller than those in the URT and vaginal microbiome simulations.

MIDASim generates read count data for each sample based on taxon relative abundances *π*_*ij*_ and library sizes *N*_*i*_. For binary-trait simulations, we considered three library-size scenarios: a non-differential scenario, where cases and controls had identical mean library sizes of 20,000, and two differential scenarios, where mean library sizes differed between cases and controls by fold changes (i.e., depth ratios) of 1.5 and 3. Specifically, a fold change of 1.5 (or 3) indicates that the mean library size of cases was 1.5 (or 3) times that of controls in the URT microbiome simulations. In contrast, for the gut microbiome simulations, a fold change of 1.5 (or 3) indicates that controls had a larger mean library size than cases. For continuous-trait simulations, only the non-differential library size scenario was considered. For each specified mean library size, the library size for each sample was drawn from a normal distribution with a standard deviation equal to one-third of the mean. Values with *N*_*i*_ *<* 2000 were resampled until *N*_*i*_ ≥ 2000.

For each simulated dataset, we first filtered out very rare taxa using the LOCOM2 filter. Table 2 compares the numbers of taxa retained under alternative filters and demonstrates that the LOCOM2 filter preserves substantially more taxa as sample size *n* increases. Consequently, the proportion of zeros also increases with *n*, e.g., rising from 61.20% at *n* = 100 to 82.41% at *n* = 1, 000 and 82.55% at *n* = 10, 000 in the URT microbiome simulations. This increasing sparsity poses growing challenges for analysis methods.

**Table 2:** Numbers of taxa passing different filters in simulation studies.

| Template dataset | Initial taxa | Filter | Taxa passing filter |  |  |
| --- | --- | --- | --- | --- | --- |
| | | | $n = 100$ | $n = 1,000$ | $n = 10,000$ |
| URT microbiome | 856 | $\min(10\%, 10)^*$ | 354.6 | 856.0 | 856.0 |
|  |  | 20%** | 177.3 | 169.1 | 170.1 |
|  |  | 10%*** | 354.6 | 329.0 | 320.4 |
| Gut microbiome<br>(CD cohort) | 2529 | $\min(10\%, 10)$ | 672.8 | 1983.6 | 2467.2 |
|  |  | 20% | 466.2 | 455.0 | 389.4 |
|  |  | 10% | 672.8 | 643.3 | 635.5 |
| Vaginal microbiome | 1146 | $\min(10\%, 10)$ | 101.1 | 375.0 | 836.6 |
|  |  | 20% | 54.4 | 54.0 | 53.6 |
|  |  | 10% | 101.1 | 99.6 | 98.8 |

Note: \*–LOCOM2 filter, which excludes taxa present in $<10\%$ of samples or $<10$ samples, whichever is smaller. \*\*–LOCOM filter, which excludes taxa present in $<20\%$ of samples. \*\*\*–Filter used by LinDA, ANCOM-BC2, and MaAsLin2, which exclude taxa present in $<10\%$ of samples. Results are based on 10 replicates of data simulated under the global null setting, i.e., $\exp(\beta) = 1$ or scale factor $s = 0$ .

We compared our proposed methods, LOCOM2 and LOCOM2-P (LOCOM2 with LOCOM-type permutation-based inference), with current state-of-the-art methods, including the original LOCOM, ANCOM-BC2 (V 2.8.1), LinDA, MaAsLin2, and MaAsLin3. For ANCOM-BC2, results were obtained after conducting pseudo-count sensitivity analysis, following the software vignette. For each method, we evaluated empirical FDR and sensitivity at the nominal FDR level of 0.2, based on 1,000 simulation replicates.

### Simulation results

Figures 3–5 present the main results, FDR and sensitivity, from simulations corresponding to the three body sites, respectively, based on the most common scenario of a binary trait with an equal case–control ratio. Each figure displayed results evaluated across three sample sizes and three library size scenarios. Among the three LOCOM-related methods, the FDR of LOCOM2 closely matched the nominal level across all scenarios, even with differential library sizes. The FDR of LOCOM2-P was sometimes conservative (Figure 3(b)) and sometimes slightly inflated (Figure 5(b)). When library sizes differed between groups, LOCOM tended to yield inflated FDR and reduced sensitivity compared with LOCOM2 and LOCOM2-P (Figure 3(a)). Moreover, at *n* = 10, 000, LOCOM and LOCOM2-P failed to produce results within one hour per replicate, whereas LOCOM2 remained scalable (∼ 10 minutes). LOCOM2-P is omitted hereafter due to the superior performance of LOCOM2.

**Figure 3:**
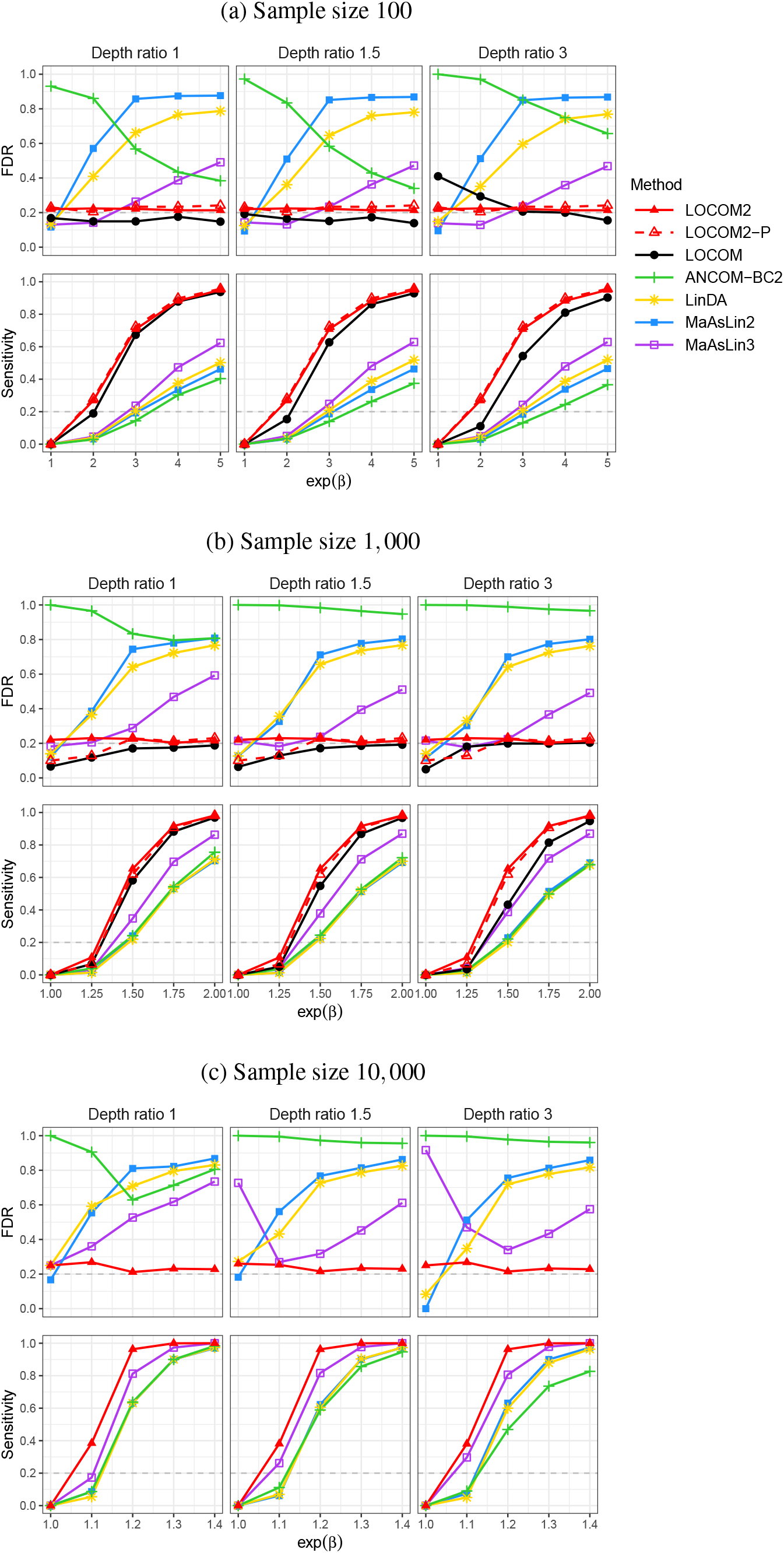
Simulation results based on the URT microbiome data template. LOCOM and LOCOM2-P are not shown for *n* = 10, 000 due to computational burden. Depth ratio is defined as the ratio of mean library sizes between cases and controls. The dashed gray line indicates the nominal FDR level of 0.2.

The other existing methods all exhibited FDR control issues in certain settings and, overall, lower sensitivity compared to LOCOM2. ANCOM-BC2 yielded highly inflated FDR when the trait effect was small or absent, and its sensitivity was generally low. The performances of LinDA, MaAsLin2, and MaAsLin3 followed a similar pattern: FDR control worsened as effect sizes increased under strong compositional effects (Figures 3 and 5), but the trend reversed when compositional effects were mild and library sizes differed (Figure 4).

**Figure 4:**
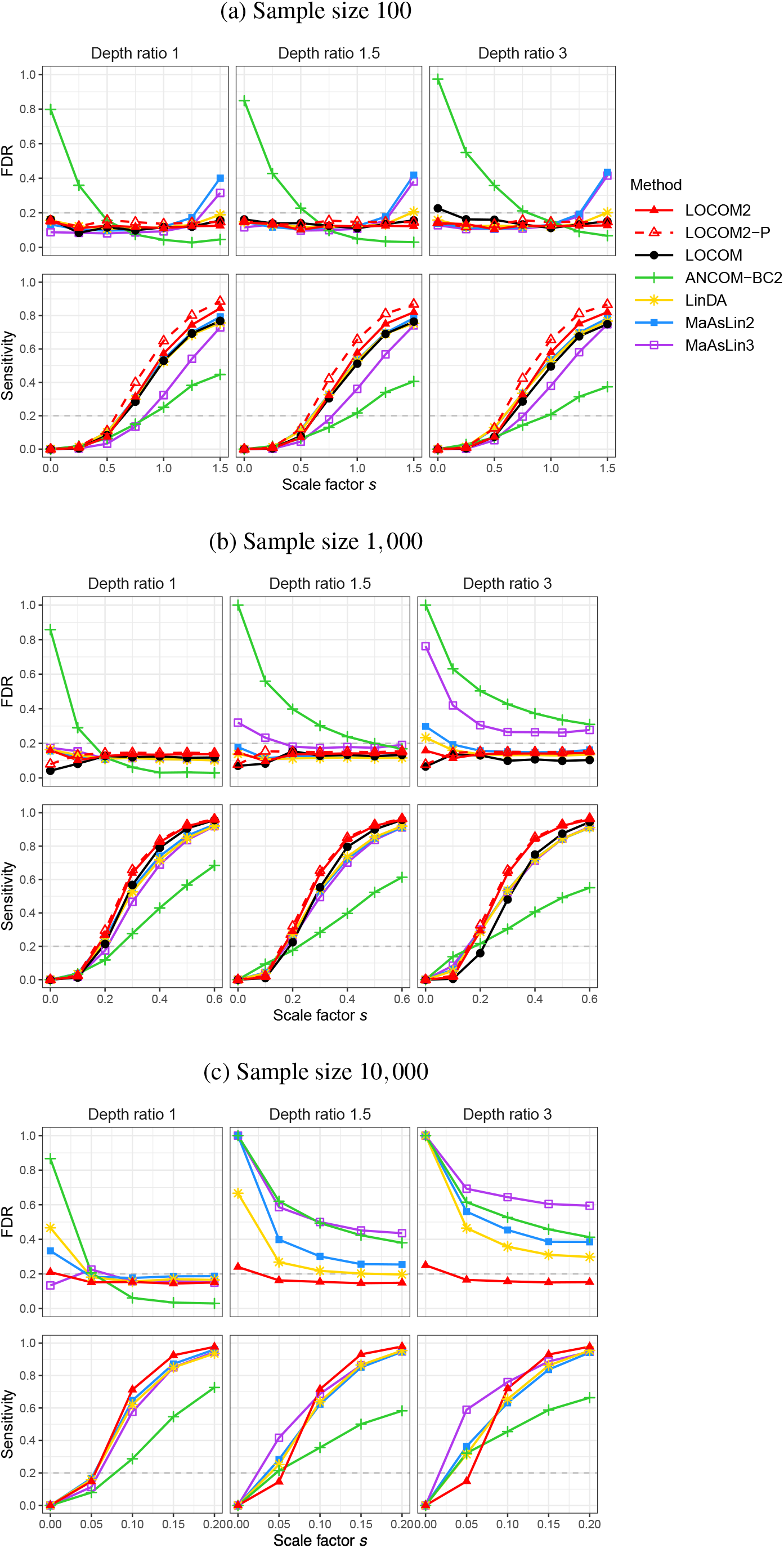
Simulation results based on the gut microbiome data template. See more information in the caption of Figure 3.

**Figure 5:**
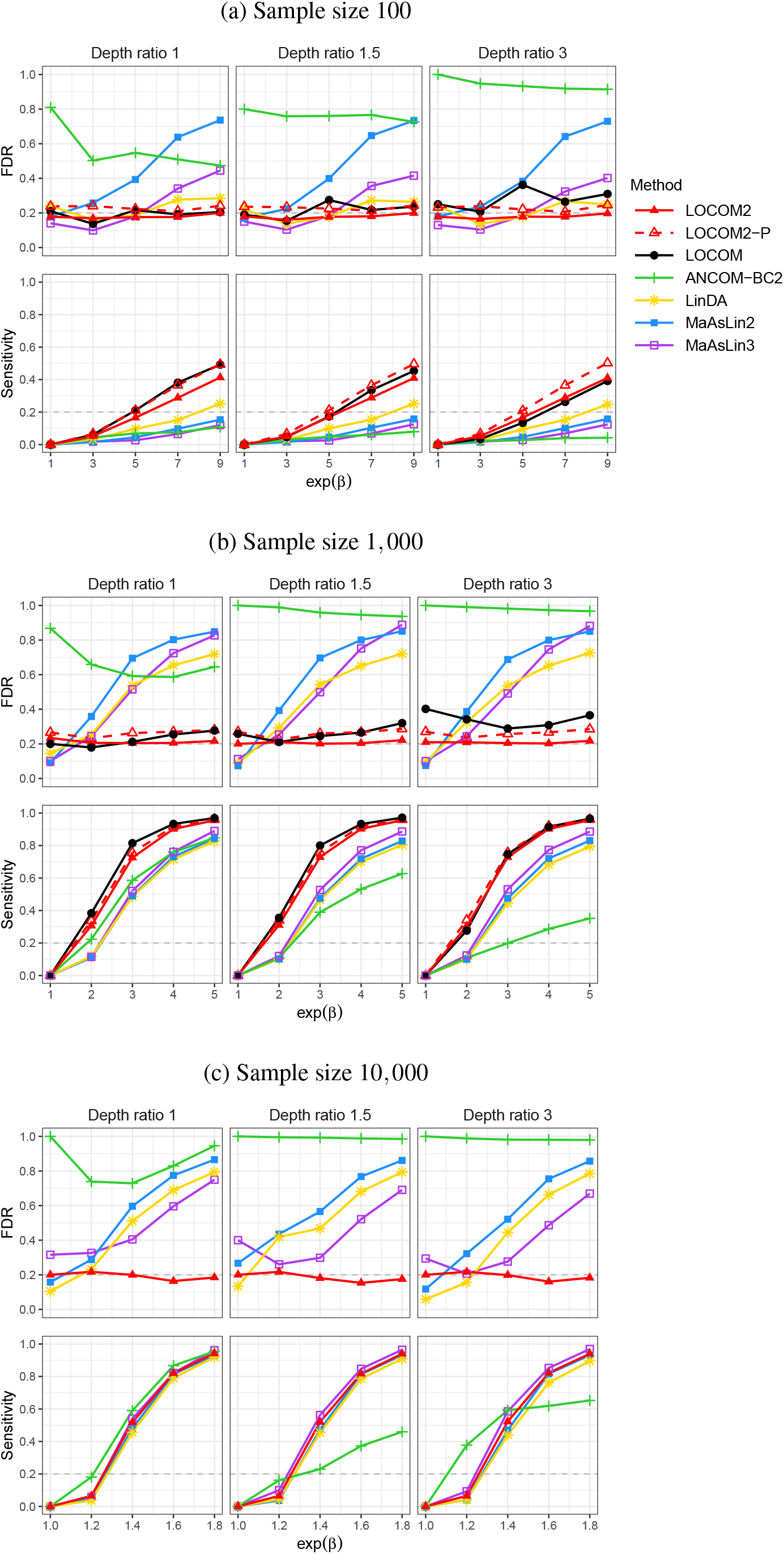
Simulation results based on the vaginal microbiome data template (from the CD cohort). See more information in the caption of Figure 3.

The results for unbalanced case–control designs are displayed in Figures 6, S6 and S7 for sample sizes of 100, 1,000, and 10,000, respectively. LOCOM2 remained robust even at extreme case-control ratios (10:90 and 90:10). LOCOM occasionally lost FDR control in these extreme scenarios, while the other competing methods continued to show FDR problems in all scenarios. The results for a continuous trait, shown in Figure 7, closely resemble those for the binary trait with non-differential library sizes.

**Figure 6:**
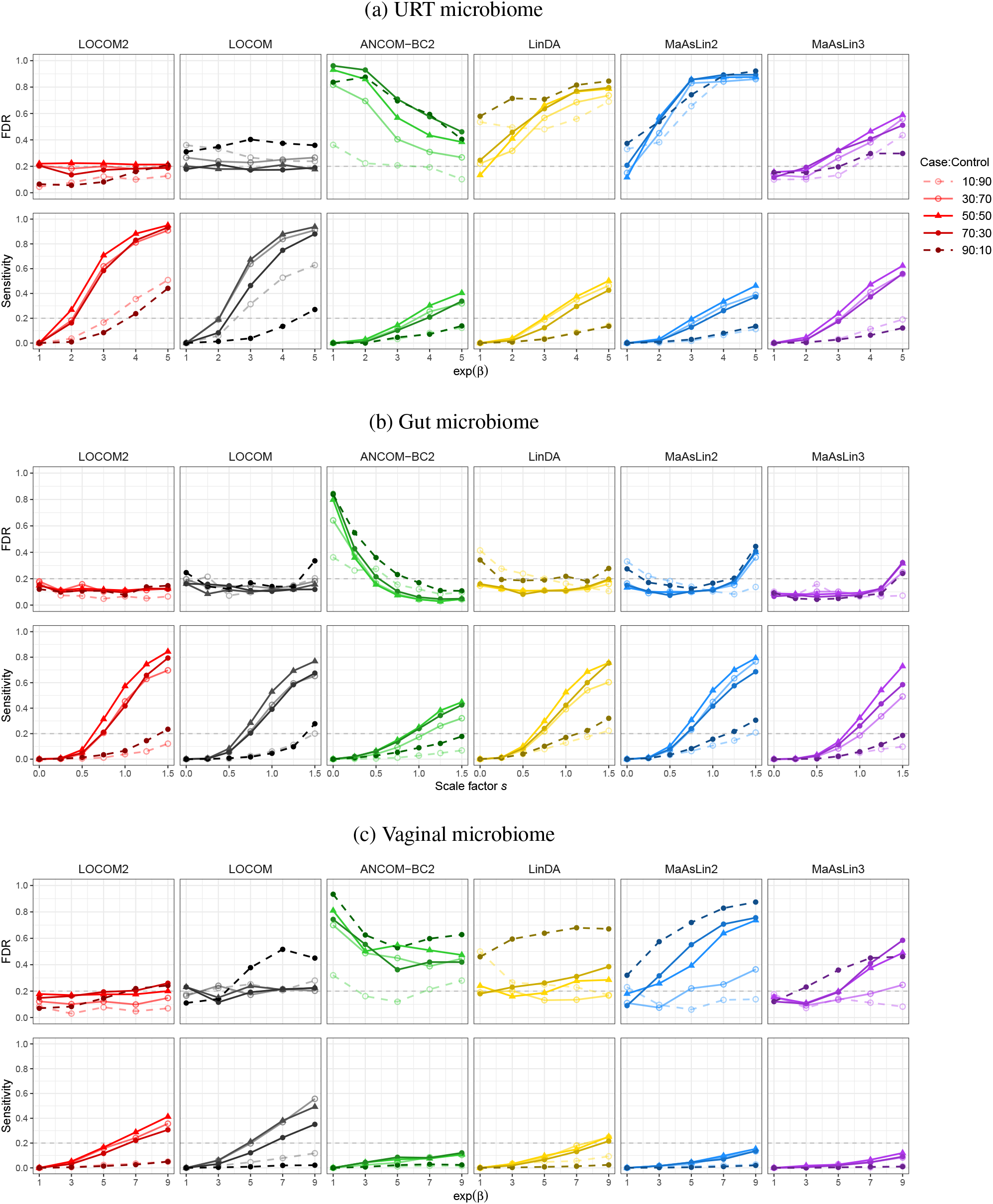
Simulation results under balanced and unbalanced case–control designs. The total sample size is *n* = 100. The dashed gray line indicates the nominal FDR level of 0.2.

**Figure 7:**
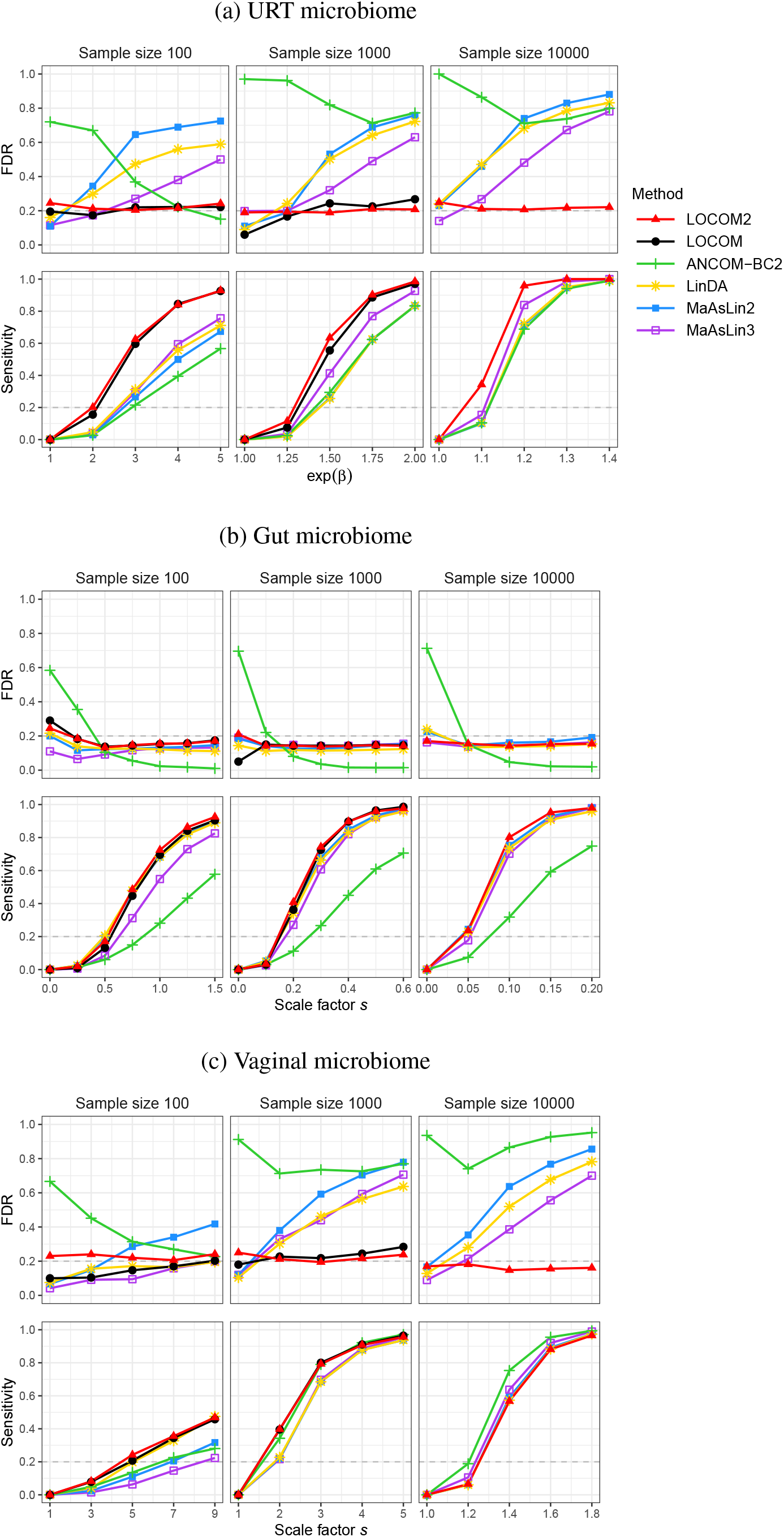
Simulation results for a continuous trait. The dashed gray line indicates the nominal FDR level of 0.2.

Figure 8 reports the computation times obtained on a 2021 MacBook Pro with an Apple M1 Pro chip and 16 GB of memory. The results show that LOCOM2 scales well to large microbiome studies: for example, analyzing a dataset with 10,000 samples required less than 12 minutes–only 1/50 of the time needed by LOCOM2-P; analyzing a dataset with 1,000 samples achieved computational efficiency comparable to ANCOM-BC2. Note that, since the permutation replicates in LOCOM2 and LOCOM2-P can be executed in parallel, they were run on four cores in parallel. In practice, many modern laptops have eight cores, which can further reduce the runtimes of LOCOM2 by approximately half compared with those reported here.

**Figure 8:**
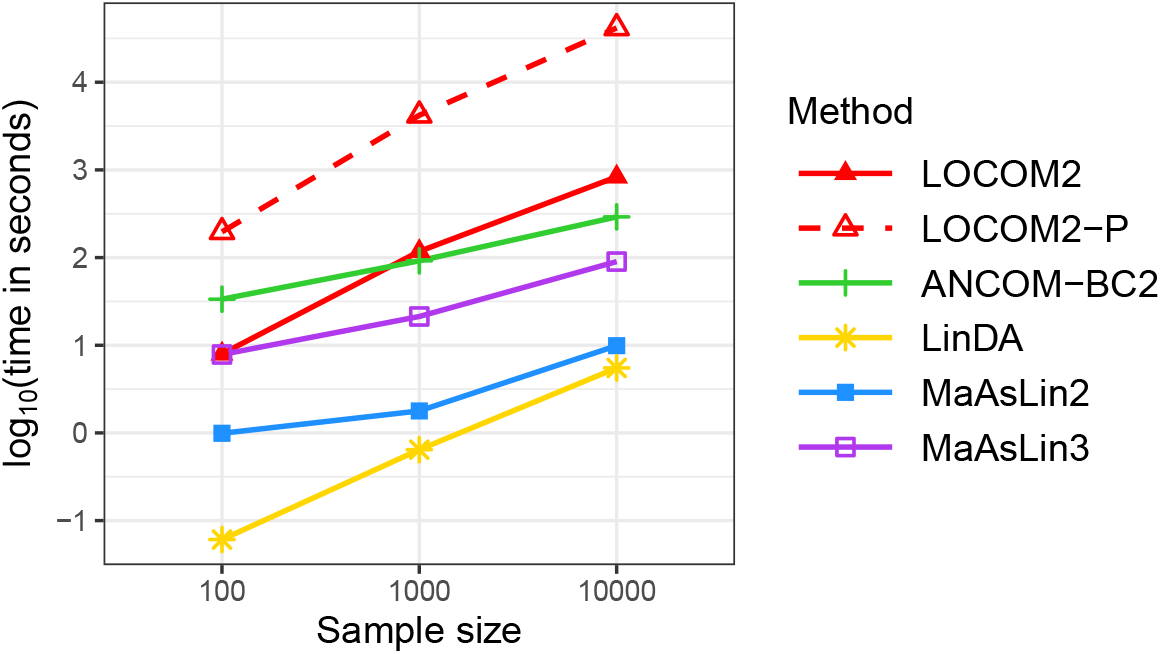
Computation time (seconds; log_10_ scale) based on 10 replicates of data simulated from the URT data template under the global null setting. LOCOM2 and LOCOM2-P were run on four cores in parallel. LOCOM is omitted as its runtime equals that of LOCOM2-P.

### Upper-respiratory-tract (URT) microbiome data

These data were originally analyzed in the LOCOM paper [12], enabling direct comparison between LOCOM and LOCOM2/LOCOM2-P. Following the same preprocessing steps as [12], we obtained 57 samples and 856 taxa. The trait of interest was smoking status, a binary variable classifying participants as 26 smokers and 31 nonsmokers. Because the proportion of males differed by smoking status, suggesting potential confounding, we adjusted for gender as a covariate. To ensure comparability with the LOCOM analysis, we applied the same filtering threshold, excluding taxa present in fewer than 20% of samples. This yielded 111 taxa for differential abundance analysis.

The results of all methods are summarized in Table 3 and Figure 9. The distributions of taxon-level *p*-values are further provided in Figure S8 to aid diagnostic checking. As previously reported [12], LOCOM detected six taxa. LOCOM2 identified eight taxa: the six found by LOCOM and two novel *Prevotella* taxa that ranked 7th and 8th by LOCOM’s *p*-values. LOCOM2 also shared five taxa with LinDA and included all three taxa identified by MaAsLin3, whereas ANCOM-BC2 did not overlap with any of the other methods. These trends align with our simulation results: LOCOM2 is slightly more powerful than LOCOM; LinDA tends to produce many false positives; MaAsLin3 has fewer errors than LinDA but lower power than LOCOM2; ANCOM-BC2 has the poorest error control and lowest power. Details of the taxa detected by LOCOM2 (i.e., taxon names, mean relative abundances, effect sizes, and *p*-values) are provided in the Supplementary File.

**Table 3:**
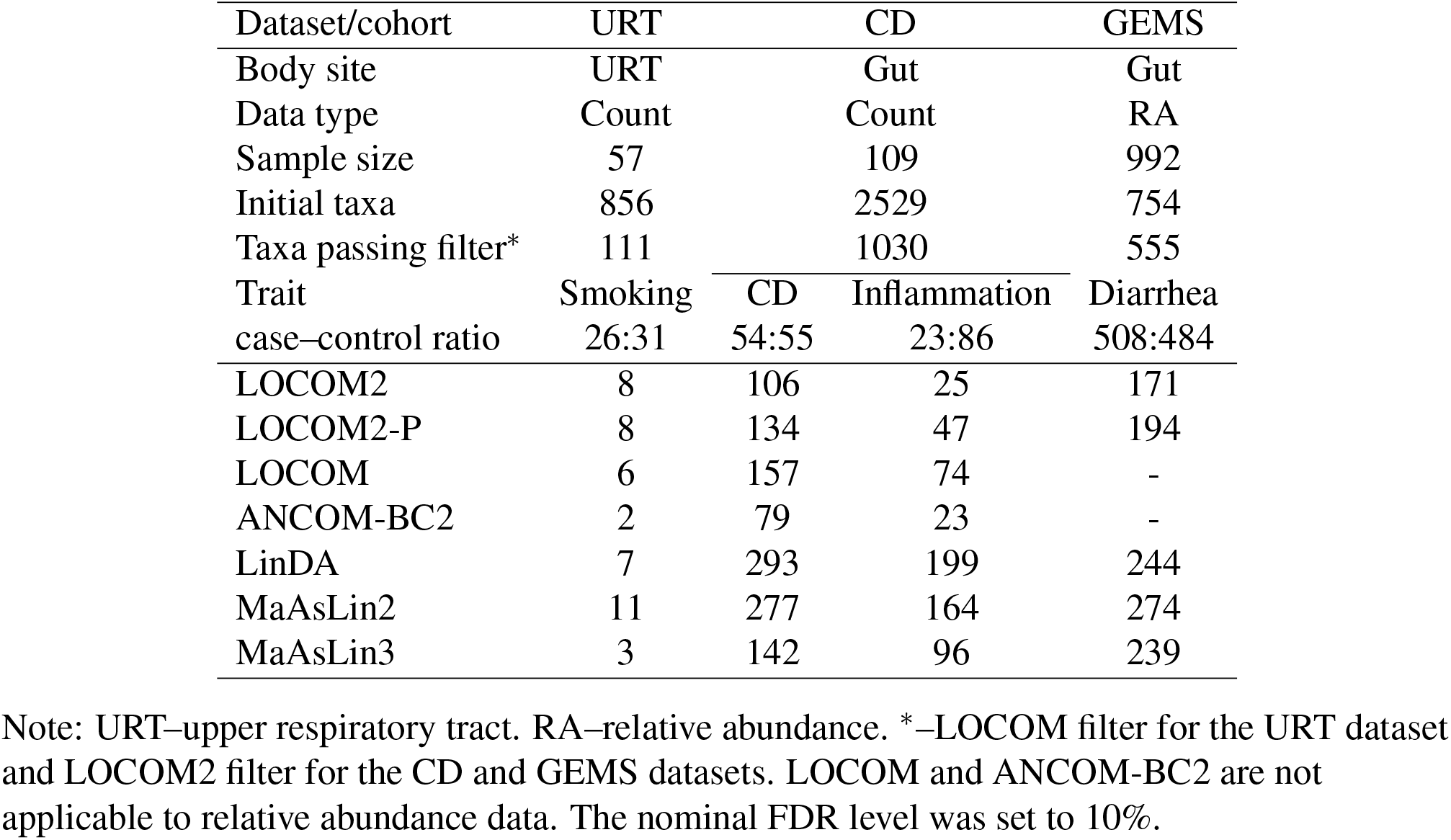
Taxa identified as differentially abundant in the real datasets.

**Figure 9:**
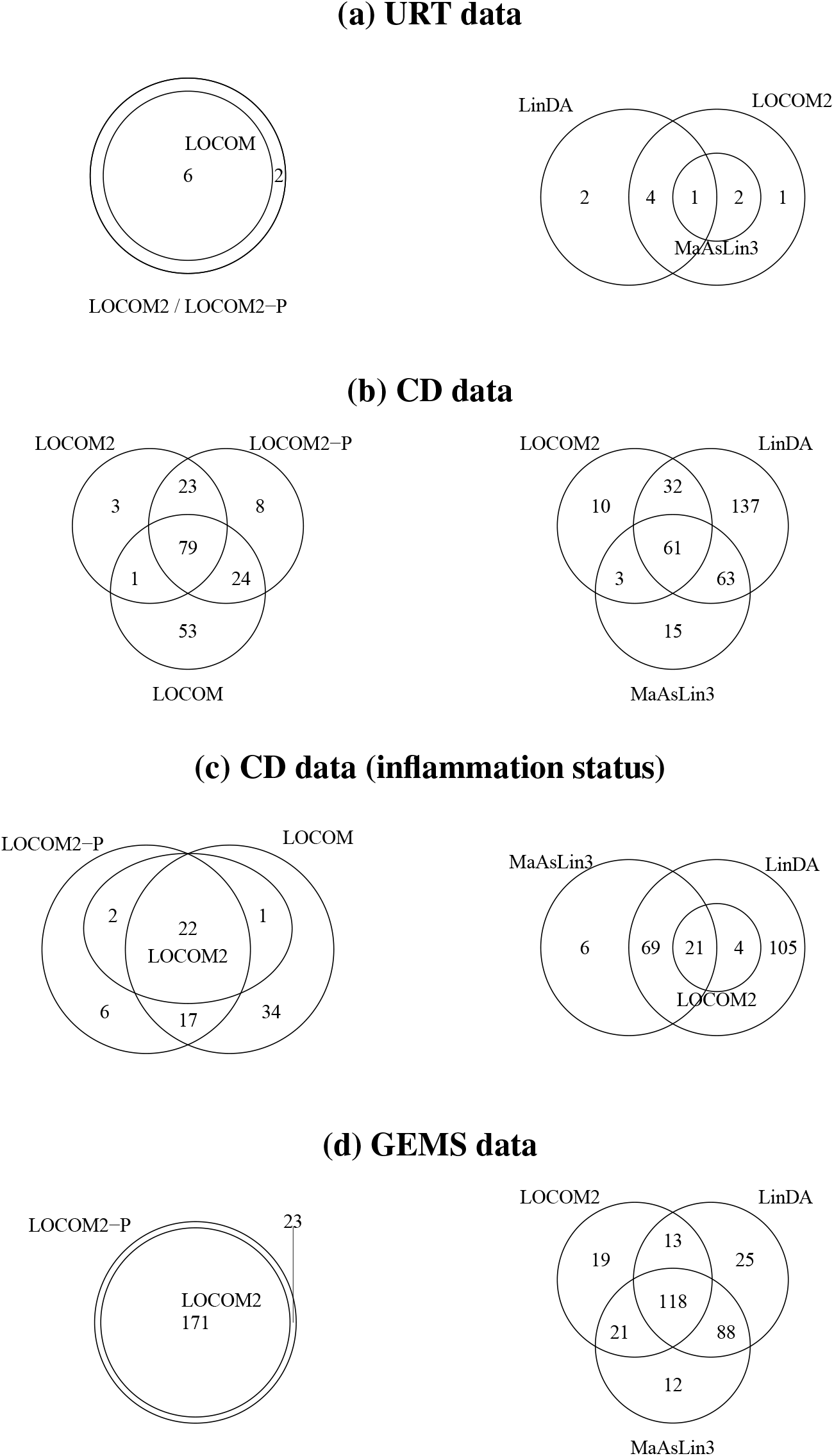
Venn diagrams of taxa identified as differentially abundant from the four analyses of three real datasets. The left panel displays results from LOCOM-related methods. The right panel compares LOCOM2 with LinDA and MaAsLin3.

### Gut microbiome data from a Crohn’s disease (CD) cohort

These data present two challenges: differential library sizes and unbalanced case–control design. It was derived from a larger inflammatory bowel disease (IBD) cohort [43] and restricted to only female subjects with CD and female controls who had not used antibiotics. After further excluding samples with library sizes below 5,000, the final dataset comprised 109 samples (54 CD cases and 55 controls) and 2,529 taxa. Applying the LOCOM2 filter yielded 1,030 taxa for differential abundance analysis.

We first analyzed the primary trait, CD status. Figure 10 shows that library sizes differed substantially between the two groups, with the mean in controls approximately 50% higher than that in cases. We then considered a secondary trait, inflammation status, yielding 23 probands with inflamation and 86 probands without inflamation, corresponding to an approximately 1:4 case–control ratio. In each analysis, age was adjusted as a potential confounder.

**Figure 10:**
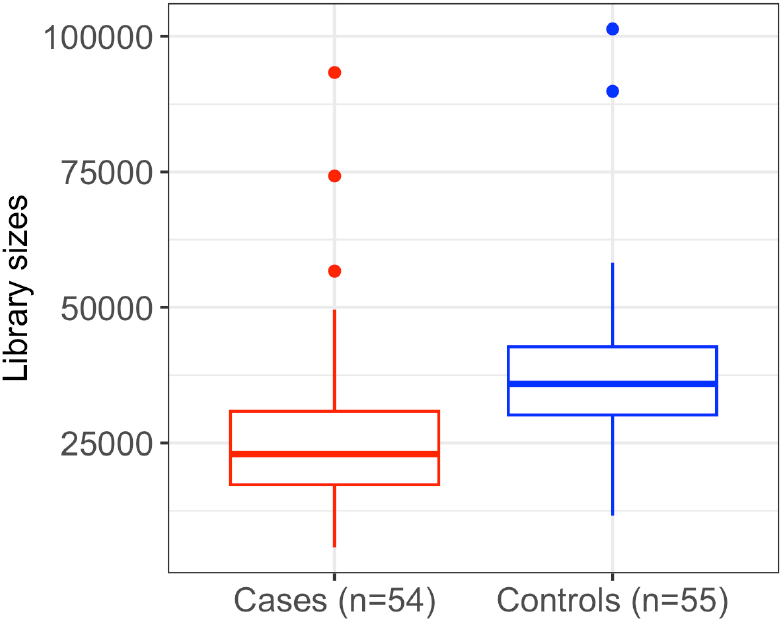
Distributions of library sizes in the gut microbiome data for CD cases and controls. The Wilcoxon rank-sum test for a location shift yielded *p <* 10^−5^.

As shown in Table 3 and Figure 9, the differential abundance results for both traits exhibit similar patterns and are consistent with our simulation findings. Taking the CD analysis as an example, LOCOM2 detected 106 taxa while LOCOM detected 157; our simulations suggest the additional detections by LOCOM are false discoveries. As expected, LOCOM2 yielded results more similar to LOCOM2-P than to LOCOM. LinDA and MaAsLin2 consistently produced the largest numbers of detections (293 and 277 taxa, respectively), followed by MaAsLin3 (142 taxa). Our simulations indicate that, in strong-signal settings, many detections by LinDA, MaAsLin2, and MaAsLin3 may be false positives. MaAsLin3 and LinDA showed greater overlap (124 taxa) with each other than with LOCOM2 (64 and 93 taxa, respectively), reflecting their shared framework of CLR-based linear regression. ANCOM-BC2 detected substantially fewer taxa overall (79 taxa), with low concordance with the other methods: only 26 detections shared with LOCOM2, 61 with LinDA, and 29 with MaAsLin3; in contrast, LOCOM2 overlapped far more with LinDA (93 taxa) and MaAsLin3 (64 taxa). The *p*-value histograms (Figure S8) reveal a clear excess of values near 1 in ANCOM-BC2’s analysis of inflammation status; in contrast, MaAsLin3’s analyses of both traits show a reduced frequency of *p*-values near 1.

### Gut microbiome data from the GEMS cohort

To demonstrate that LOCOM2 scales efficiently to large-scale microbiome data and accommodates relative-abundance inputs, we analyzed the pediatric subset of the Global Enteric Multicenter Study (GEMS) case–control cohort [44] that included data from 992 children under five years of age. These children comprise 508 moderate–severe diarrhea cases and 484 diarrhea-free controls, with covariates for age (0–59 months) and recruitment site (Bangladesh *n* = 206, Gambia *n* = 269, Kenya *n* = 305, Mali *n* = 212). Relative abundance data were initially available for 754 taxa, of which 555 remained after applying the LOCOM2 filter. By contrast, the original LOCOM filter (20% prevalence) and the ANCOM-BC2 filter (10% prevalence) would have retained only 110 and 188 taxa, respectively. We assessed the association between diarrhea status and 555 taxa in the 992 children, adjusting for age and recruitment site.

Because only relative abundance data were available, LOCOM and ANCOM-BC2 were not applied. As shown in Table 3 and Figure 9, LOCOM2 detected 171 taxa, whereas LOCOM2-P detected 194; some of the 23 additional taxa identified by LOCOM2-P but not by LOCOM2 had relatively large *p*-values under LOCOM2. LinDA and MaAsLin2 again produced the largest numbers of detections (244 and 274 taxa, respectively), followed by MaAsLin3 (239 taxa). MaAsLin3 and LinDA showed greater overlap (206 taxa) with each other than either did with LOCOM2 (139 and 131 taxa, respectively).

## Discussion

The reproducibility crisis of microbiome research is largely attributable to the inability to use internal standards to normalize microbial count data. This is a consequence of the findings of [3], who used data from model microbial communities to show that each taxon was subject to its own “bias factor” that alters the measured counts in a multiplicative fashion. Because each taxon has its own bias factor, we would require an internal standard for every taxon we observe. Because this is not feasible, the only currently-available option is to only use analysis methods that automatically compensate for these biases. Worse, because bias factors depend on both the extraction and sequencing protocols used, the same true relative abundance in two studies may appear to differ, even when log-ratios are used.

Addressing the replicability crisis in microbiome research requires differential abundance analysis methods that reliably control error rates and produce stable, reproducible findings across independent studies. By maintaining rigorous FDR control across diverse settings, including largescale datasets, differential library sizes, unbalanced case–control designs, and relative-abundance–only inputs, LOCOM2 reduces the risk of spurious discoveries. At the same time, its improved sensitivity relative to existing approaches enhances the efficiency of detecting true biological signals. Together, these properties make LOCOM2 a powerful method to advance reproducible microbiome research.

LOCOM2 is currently limited to tests having only one degree of freedom. The test statistic in (4) can easily be generalized to multiple degrees of freedom; the research question is what multivariate median should be used to center the (now multivariate) values of 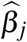 when forming *T*_*j*_. We hope to address this question soon.

This study has two limitations. First, we did not conduct simulations based on the GEMS dataset, as we had already performed simulations using the CD dataset, which also pertains to the gut microbiome. Second, because the MOMS-PI data (obtained from the R package HMP2Data) lack an associated trait, we did not conduct a real-data analysis for this dataset.

LOCOM2 shows substantially improved performance analyzing rare taxa compared to LOCOM, allowing a much less stringent filtering criterion. In practice, however, we may not always want such a liberal filtering criterion, because correcting for multiple testing can impose a harsh penalty on statistical power. For example, when the LOCOM2 filter was applied to the URT dataset, LOCOM2 detected only four taxa rather than the eight detected when the filtering criterion for LOCOM was used. Nevertheless, our results indicate that LOCOM2 is well positioned to reliably analyze data from rare taxa.

In its output, LOCOM2 provides, for each taxon, an effect size estimate 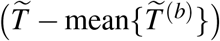 together with its standard error estimate 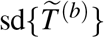. These quantities enable *meta-analysis*, which combines results across multiple independent microbiome studies to obtain an overall estimate, using the standard inverse-variance-weighted method. As microbiome studies continue to grow and large consortia [45] start to emerge, this feature is essential towards reproducible and generalizable microbiome discoveries.

