## Supplemental materials for "LOCOM2: Robust Differential Abundance Analysis for Microbiome Data"

### Supplementary Materials

#### Adjusted estimating equation

We adopt the general framework of adjusted estimating equations proposed by Kosmidis and Lunardon (K-L) [36]. In this supplement, we suppress the index $j$ for notional simplicity, with the understanding that the procedure is applied to each taxon $j$. We redefine $Z_i$ to denote the full covariate vector of length $K$, including the intercept term 1. We also redefine $\theta$ to denote the full parameter vector, including the intercept $\eta$ and any nuisance parameters $\phi$. Under these definitions, the logistic model in (2) can be rewritten as

$$\log\left\{\mu_{ij}/(1-\mu_{ij})\right\} = \theta^{\mathrm{T}} Z_i \ .$$

K-L derived an analogue of the Firth correction for estimating equations. However, unlike the likelihood-based Firth correction, this bias-reduction approach does not guarantee a finite estimator of $\theta$ when there is separation in the data. To address this issue, K-L proposed augmenting the estimating equations with an additional Jeffreys-type penalty to ensure finiteness [37]. Following this approach, our adjusted estimating equation is given by

$$U(\theta) + A(\theta) + G(\theta) = 0 \ , \tag{S1}$$

where $U(\theta)$ denotes the estimating equation for logistic regression, $A(\theta)$ is the K-L bias-reduction adjustment, and $G(\theta)$ is the Jeffreys-type penalty term. We describe each of these components below. Notably, to implement the K-L approach, we must choose the scaling such that $U(\theta) = O(n)$, $A(\theta) = O(1)$ and $G(\theta) = O(1/n)$.

The first term, $U(\theta)$, corresponds to the standard logistic regression estimating equation. To ensure proper scaling, we rewrite and rescale $U(\theta)$ in (3) with $\omega_i = 1$ as

$$U(\theta) = n \sum_{i=1}^{n} \widetilde{\omega}_i \left( \frac{\widehat{p}_i}{\widehat{m}_i} - \mu_i \right) Z_i \ ,$$

where $\widehat{m}_i = \widehat{p}_i + \widehat{p}_J$ (recall that $J$ is the reference taxon) and $\widetilde{\omega}_i = \widehat{m}_i / \sum_{i'} \widehat{m}_{i'}$, so that $\sum_i \widetilde{\omega}_i = 1$.

Under this formulation, $U(\theta) = o(n)$. We further define $U_1(\theta) = n^{-1}U(\theta)$ as the contribution of a single observation (e.g., observation 1) to $U(\theta)$.

The second term, $A(\theta)$, represents the K-L bias-reduction adjustment, which reduces the asymptotic bias of estimators obtained from (asymptotically unbiased) estimating equations. The $r$-th component of the $K$-vector $A(\theta)$ is defined as

$$A_r(\theta) = -\operatorname{Tr}\left\{J(\theta)^{-1} d_r(\theta)\right\} - \frac{1}{2}\operatorname{Tr}\left\{J(\theta)^{-1} e(\theta)\left[J(\theta)^{-1}\right]^{\mathrm{T}} u_r(\theta)\right\} ,$$

where $J(\theta)$, $d_r(\theta)$, $e(\theta)$, and $u_r(\theta)$ are arrays with elements given by

$$J_{kl}(\theta) = -\mathrm{E}\left(\frac{\partial U_{1,k}}{\partial \theta_l}\right) ,$$

$$d_{rkl}(\theta) = \mathrm{E}\left[\left(\frac{\partial U_{1,r}}{\partial \theta_k}\right) U_{1,l}\right] ,$$

$$e_{kl}(\theta) = \mathrm{E}\left(U_{1,k}U_{1,l}\right) ,$$

and

$$u_{rkl}(\theta) = \mathrm{E}\left(\frac{\partial U_{1,r}}{\partial \theta_k \partial \theta_l}\right) .$$

These quantities can be replaced by their empirical estimates:

$$\widehat{J}_{kl}(\theta) = \sum_{i=1}^{n} \widetilde{\omega}_i \mu_i (1-\mu_i) Z_{ik} Z_{il} ,$$

$$\widehat{d}_{rkl} = \sum_{i=1}^{n} \widetilde{\omega}_i \left(\frac{\widehat{p}_i}{\widehat{m}_i} - \mu_i\right) \mu_i (1-\mu_i) Z_{ir} Z_{ik} Z_{il} ,$$

$$\widehat{e}_{kl}(\theta) = \sum_{i=1}^{n} \widetilde{\omega}_i \left(\frac{\widehat{p}_i}{\widehat{m}_i} - \mu_i\right)^2 Z_{ik} Z_{il} ,$$

and

$$\widehat{u}_{rkl} = -\sum_{i=1}^{n} \widetilde{\omega}_i \mu_i (1-\mu_i)(1-2\mu_i) Z_{ir} Z_{ik} Z_{il} .$$

With these definitions, it is clear that $A(\theta) = o(1)$, as required.

The third term, $G(\theta)$, corresponds to the Jeffreys-type penalty:

$$G(\theta) = (2n)^{-1} \frac{\partial \log |J(\theta)|}{\partial \theta}.$$

The empirical estimate of $J(\theta)$ can be expressed in matrix form as

$$\widehat{J}(\theta) = \mathbb{Z}^{\mathrm{T}} \Omega \mathbb{Z} \, ,$$

where $\mathbb{Z} = (Z_1, Z_2, \ldots, Z_n)^{\mathrm{T}}$ and $\Omega = \mathrm{diag}\{\widetilde{\omega}_i \mu_i (1 - \mu_i)\}$. Then, standard matrix calculus yields

$$G(\theta) = \frac{1}{2n} \sum_{i=1}^{n} h_{ii} (1 - 2\mu_i) Z_i \, ,$$

where $h_{ii}$ denotes the $i$-th diagonal element of the hat matrix $H = \Omega^{\frac{1}{2}} \mathbb{Z} \left( \mathbb{Z}^T \Omega \mathbb{Z} \right)^{-1} \mathbb{Z}^T \Omega^{\frac{1}{2}}$. Since $\mathrm{Tr}(H) = K = o(1)$, each diagonal element $h_{ii}$ is of order $1/n$, implying that $G(\theta) = o(1/n)$, as required.

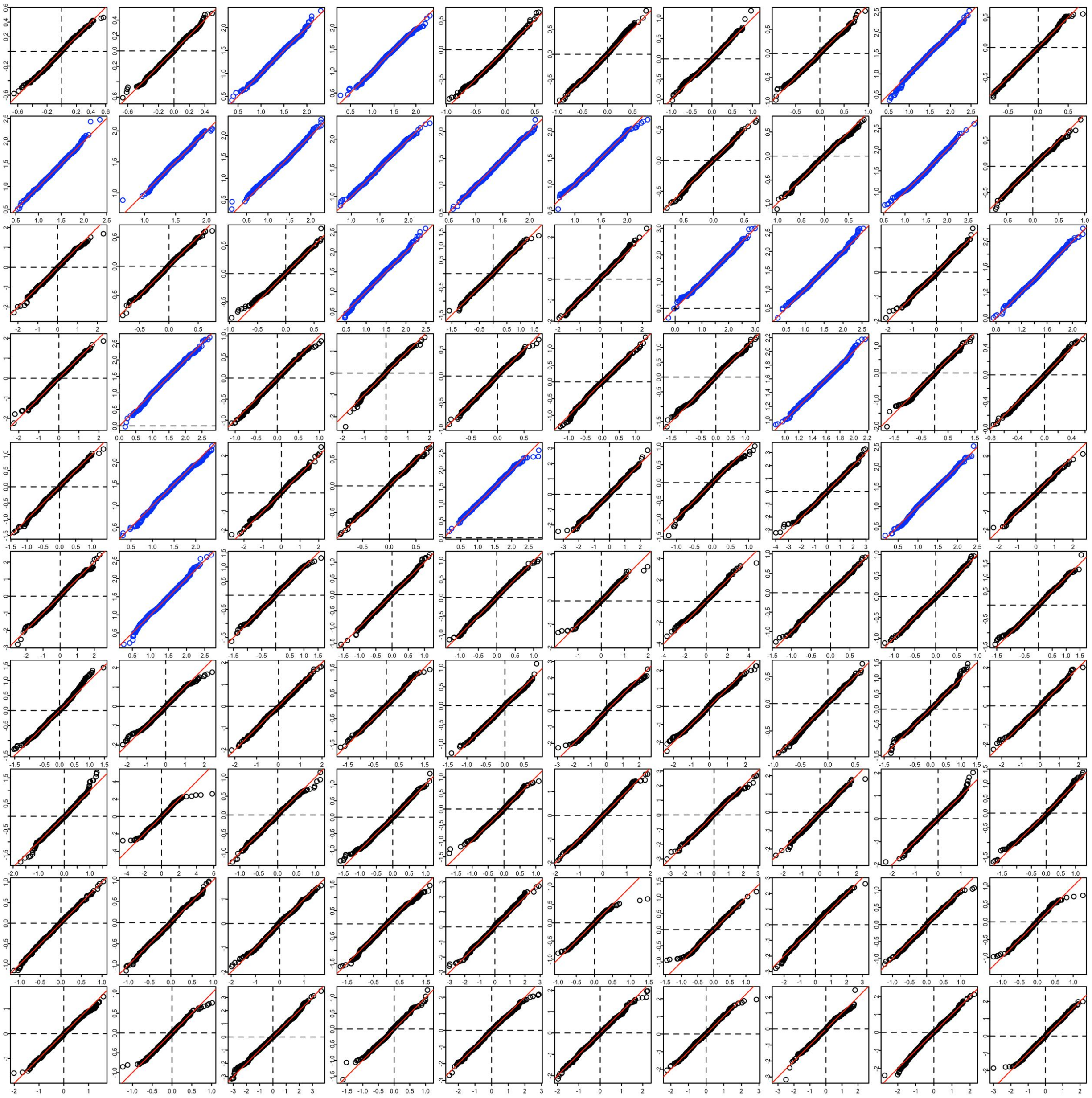

Figure S1: Quantile–quantile (QQ)-plots of the $T$ statistics from 1,000 simulation replicates against normal distributions (using the per-taxon mean and variance of the $T$ statistics). Results are shown for the top 100 most abundant taxa (out of 193 total; blue denotes differentially abundant taxa and black denotes null taxa), based on simulations of the URT microbiome data with $n = 100$, $\exp(\beta) = 5$, depth ratio $= 1$, and an equal case–control ratio. Plots are arranged from left to right and top to bottom in descending order of mean relative abundance; each plot shows the 45° reference line (solid red) and zero reference lines on both axes (black dashed).

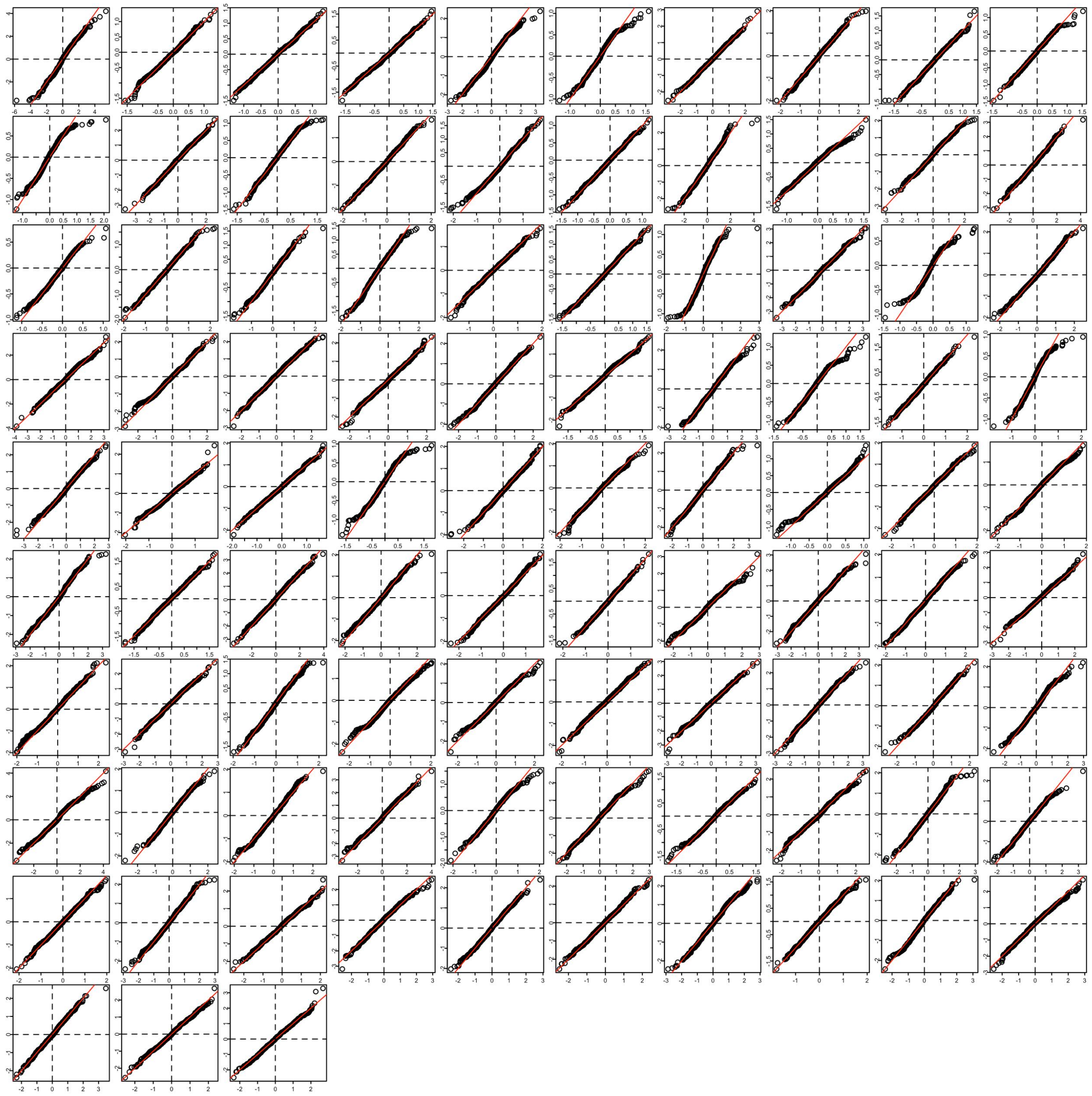

Figure S2: QQ-plots for the remaining 93 less abundant taxa. See more information in the caption of Figure S1.

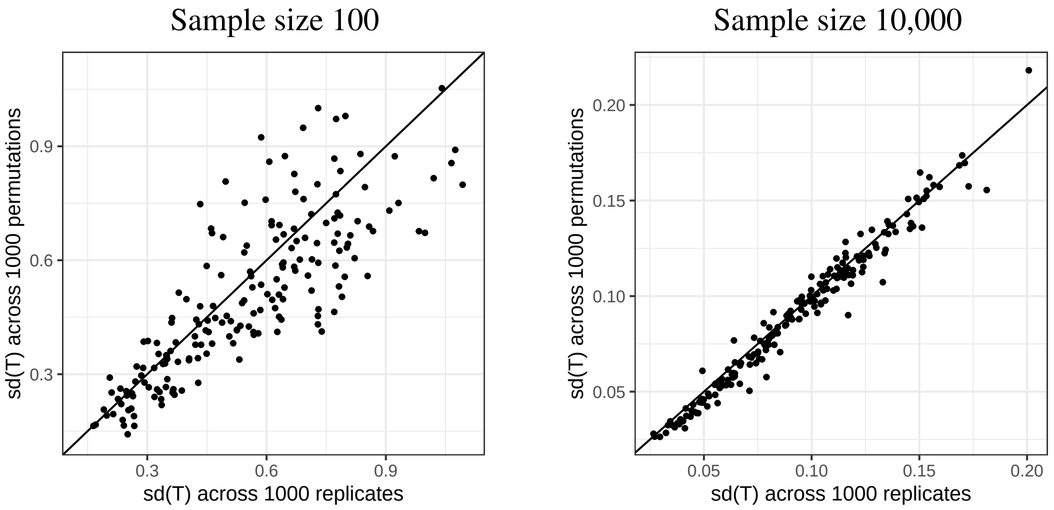


Figure S3: Comparison of the standard deviation of the $T$ statistics from 1,000 simulation replicates (x-axis) and from 1,000 permutations of a single replicate (y-axis), based on simulations of the URT microbiome data with an equal case–control ratio and depth ratio = 1. The left panel corresponds to $n = 100$ and $\exp(\beta) = 5$; the right panel corresponds to $n = 10,000$ and $\exp(\beta) = 1.4$.

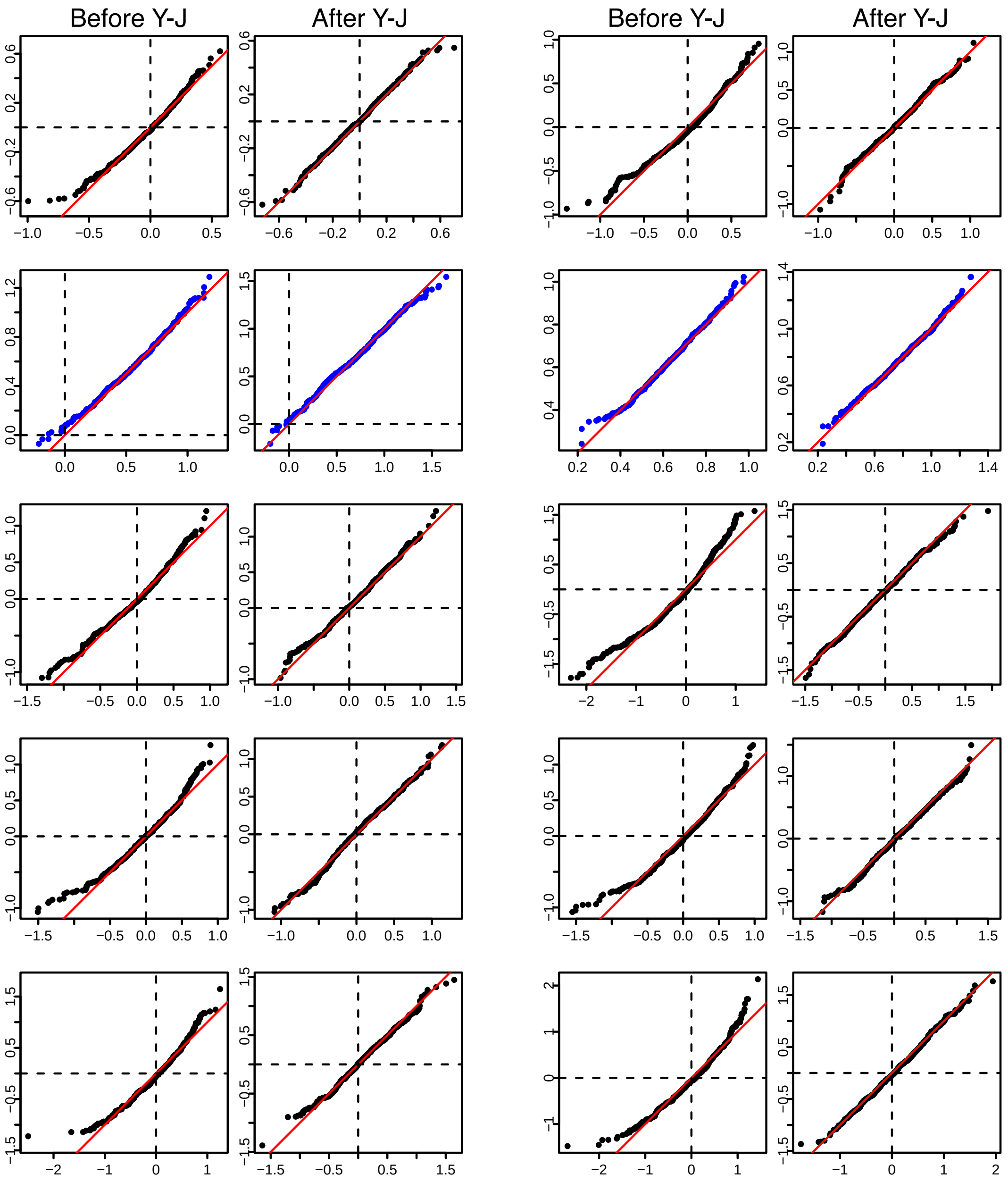


Figure S4: Comparison of the $T$ statistics before and after Yeo–Johnson (Y-J) transformation. QQ-plots are shown for ten taxa with $\widehat{\lambda} > 1.6$, based on 1,000 simulation replicates of the URT microbiome data with $n = 1{,}000$, $\exp(\beta) = 2$, depth ratio $= 1$, and case–control ratio of 1:9. See more information in the caption of Figure S1.

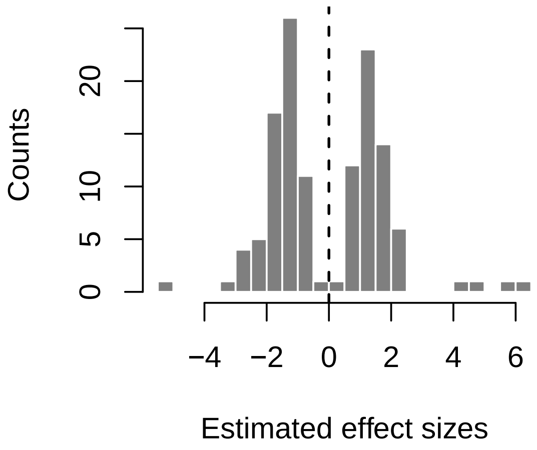


Figure S5: Histogram of estimated effect sizes for 106 taxa identified as differentially abundant in the gut microbiome data from the CD cohort.

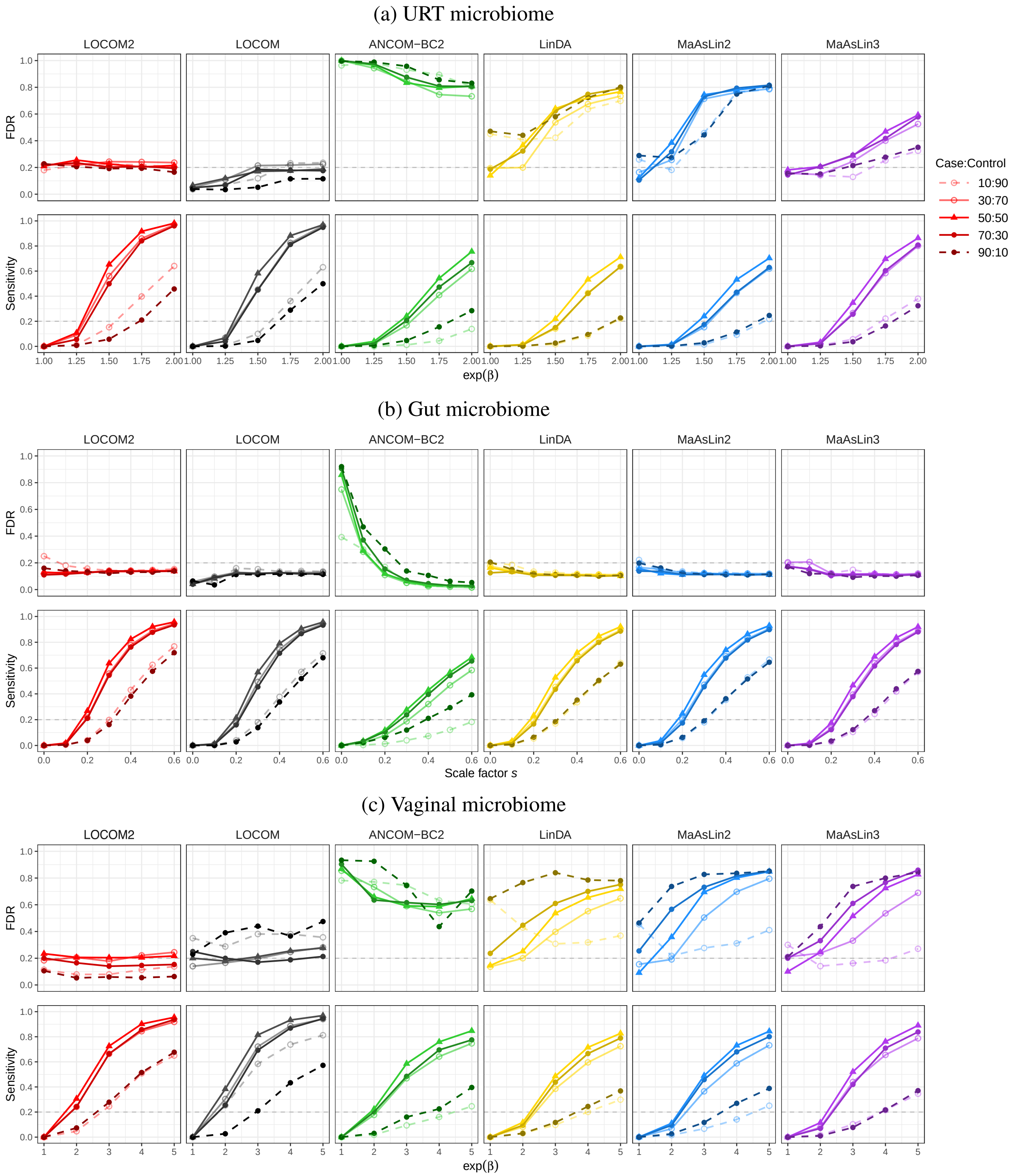


Figure S6: Simulation results under balanced and unbalanced case–control ratios with a sample size of $n = 1,000$. The dashed gray line indicates the nominal FDR level of 0.2.

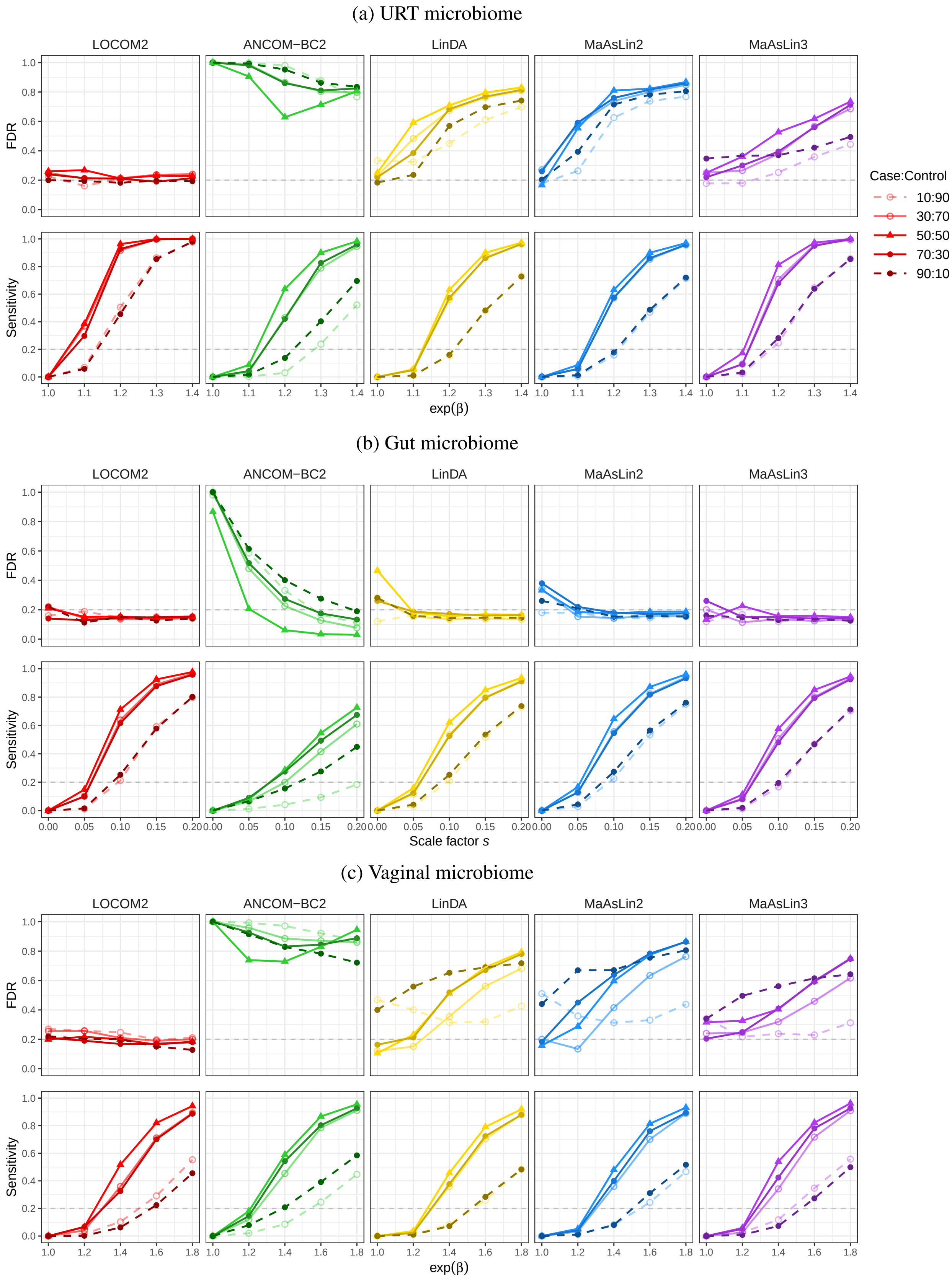


Figure S7: Simulation results under balanced and unbalanced case–control ratios with a sample size of $n = 10,000$. The dashed gray line indicates the nominal FDR level of 0.2.

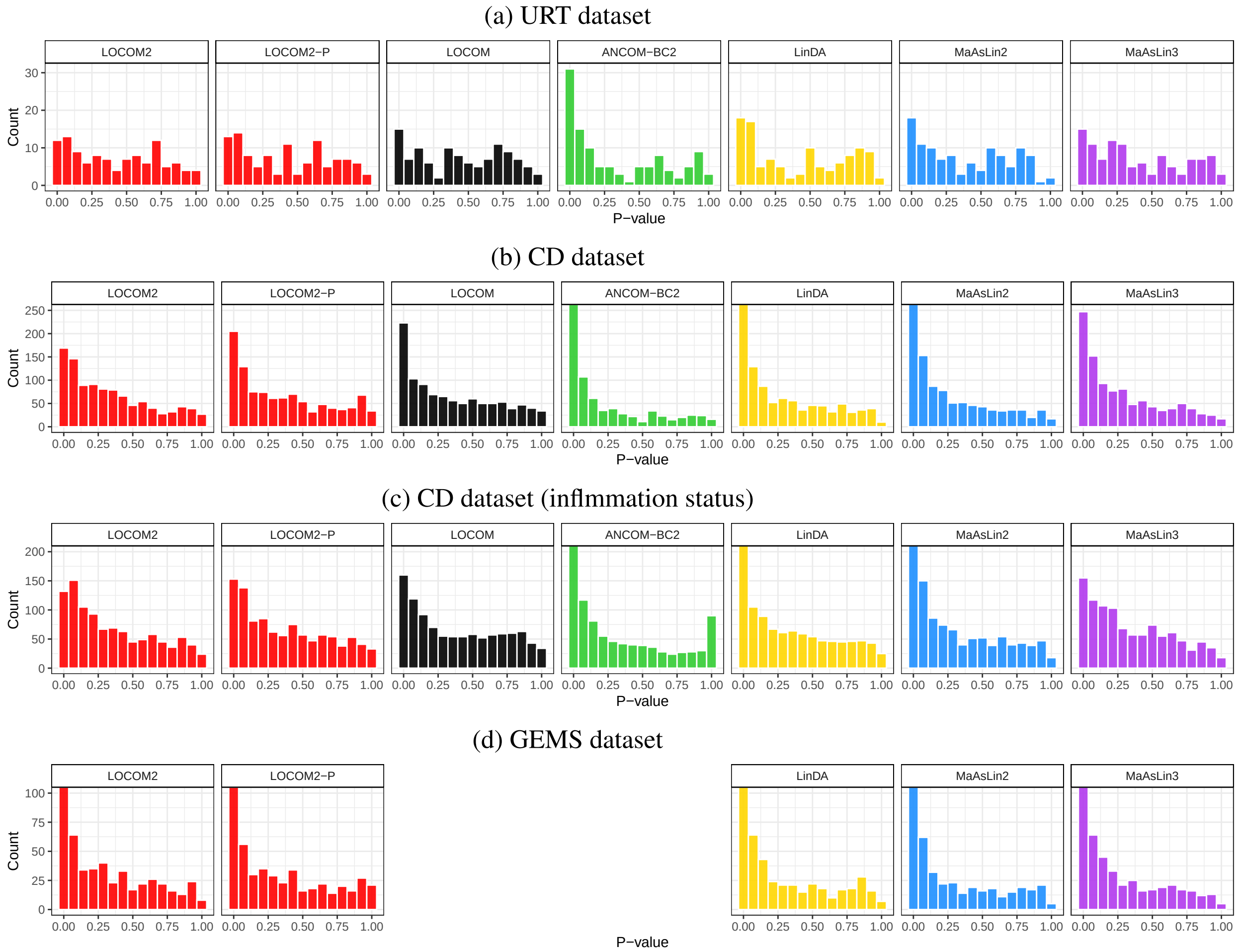


Figure S8: Histograms of taxon-level $p$-values in analyses of real datasets. LOCOM and ANCOM-BC2 were not applied to the GEMS dataset as only relative abundance data were available.
